# Subcompartmentalization and pseudo-division of model protocells

**DOI:** 10.1101/2020.06.07.138479

**Authors:** Karolina Spustova, Elif Senem Köksal, Alar Ainla, Irep Gözen

## Abstract

Membrane enclosed intracellular compartments have been exclusively associated with the eukaryotes, represented by the highly compartmentalized last eukaryotic common ancestor. Recent evidence showing the presence of membranous compartments with specific functions in archaea and bacteria makes it conceivable that the last universal common ancestor and its hypothetical precursor, the protocell, could have exhibited compartmentalization. To our knowledge, there are no experimental studies yet that have tested this hypothesis. We report on an autonomous subcompartmentalization mechanism for protocells which results in the transformation of initial subcompartments to daughter protocells. The process is solely determined by the fundamental materials properties and interfacial events, and do not require biological machinery or chemical energy supply. In the light of our findings, we propose that similar events could have taken place under early Earth conditions, leading to the development of compartmentalized cells and potentially, primitive division.

---

The origin of the eukaryotic cell is closely associated with the development of subcompartments, which create specific micro-environments to spatially or temporally regulate biochemical reactions, simultaneously. Until recently, cellular compartmentalization was associated solely with eukaryotic systems. Recent evidence shows that membrane-enclosed compartments also exist in other domains of life, such as in Archaea and Bacteria^1,2^. Archaea, for example, have acidocalcisomes, the membrane enclosed electron-dense granular organelles rich in calcium and phosphate, which is crucial for osmoregulation and calcium homeostasis^3^. In Cyanobacteria, membrane-bound thylakoids^4^, have been identified as compartments in which the light-dependent reactions of photosynthesis take place.

Despite the differences between eukaryotic and prokaryotic compartments in terms of structural and functional complexity, the presence of membranous compartments in Procaryota^1^ establishes a possibility of compartments having existed in protocells, and being evolutionarily conserved. There is essentially no experimental material, however, on how compartments could have consistently emerged from membranes in a prebiotic environment lacking membrane-shaping and -stabilizing proteins.

Membrane-less laboratory models of cytoplasmic suborganization have been developed inside synthetic cells, *i.e*. giant unilamellar vesicles^5^. These models require moderately elaborate chemical systems. A few examples of the utilized materials and mechanisms to induce compartment formation involve thermo-responsive hydrogels^6^, pH driven protein (human serum albumin) localization^7^ or a poly(ethylene glycol)-dextran aqueous two-phase system^8^. One study which report on the membrane-based multi-vesiculation inside amphiphilic compartments, has employed protein-ligand couples to induce this behavior, *i.e*. biotin-avidin conjugates^9^. Membrane-enveloped subcompartment formation in biological systems is therefore considered to be ultimately dependent on the genetic make-up of the cells^10^.

In this study, we have investigated a protocell subcompartmentalization model on solid surfaces. The reported mechanism relies on the self-assembly and shape transformation abilities of biosurfactant bilayer membranes, and is largely governed by changes in the membrane-surface adhesion. Model protocells in form of giant unilamellar compartments are initially adhered to a solid substrate via Ca^2+^ anchors, but the adhesion is subsequently reversed upon depletion of Ca^2+^. Small membrane regions released from the surface transform into small unilamellar subcompartments inside the model protocell. They remain partially adhered to the surface and continue growing, consuming the membrane material of the primary container. The subcompartments can take up solutes from the external environment and exchange them with the primary volume. Our experiments and finite element simulations point to a parallel uptake mechanism of solutes via transient pores in the protocell membrane. The enveloping membrane can eventually rupture to release the subcompartments as independent daughter vesicles.

Previous studies indicate that surfaces might have played a role in catalysis and synthesis of compounds relevant to prebiotic chemistry, *e.g*. enhanced polymerization of aminoacids on mineral surfaces^11^, or the synthesis of prebiotic peptides^12^ and RNA^13^. Our work shows that the surface-assisted subcompartmentalization of protocells is a feasible recurring process and, due to the minimal requirements, could have potentially occurred under early Earth conditions. The newly established pathway to daughter vesicles, although relatively unsophisticated, indicates that the role of materials properties-driven transformation processes of surfactant membranes in the development of primitive protocells, as well as the enabling function of surfaces, might have been underestimated.

## Results

### Spontaneous compartmentalization

**Fig. 1a** is a 3D reconstruction of the subcompartments that have spontaneously formed inside a substrate-adherent model protocell. The dome-shaped lipid container of approximately 50 µm in diameter is adhered on an Al_2_O_3_ surface where its distal membrane envelopes tens of subcompartments of an average size of 5 µm formed on the surface. **Fig. 1b** shows an xy cross sectional view recorded 4 µm above the surface. **Fig. 1c-f** contain confocal micrographs of the compartmentalization process each step of which is explained in schematics in **Fig. 1g-j**. The model protocells were generated by the dehydration–rehydration method^14^; and transferred onto the solid substrates in presence of Ca^2+^ as adhesion promoter^15,16^(**Fig. 1c** and **1g**). Subsequently, by means of an automatic pipette, the ambient buffer was exchanged with a Ca^2+^chelator-containing buffer, leading to the gradual removal of Ca^2+^ from the membrane-surface interface (**Fig. 1d** and **1h**). The basal membrane, *i.e*. the adhered portion of the giant lipid container to the surface, de-wets the surface, bulges at various locations (**Fig. 1e** and **1i**) and transform within ∼40 min into spherical subcompartments (**Fig. 1f** and **1j**). The model protocells in **Fig. 1c-d** are isolated. The elevated background intensity should not be mistaken for a lipid film. The substrate is composed of Al which reflects light during the imaging. A lipid reservoir, *i.e*. multilamellar vesicle (MLV), is attached to the model protocell in **Fig. 1c**.

**Figure 1.**
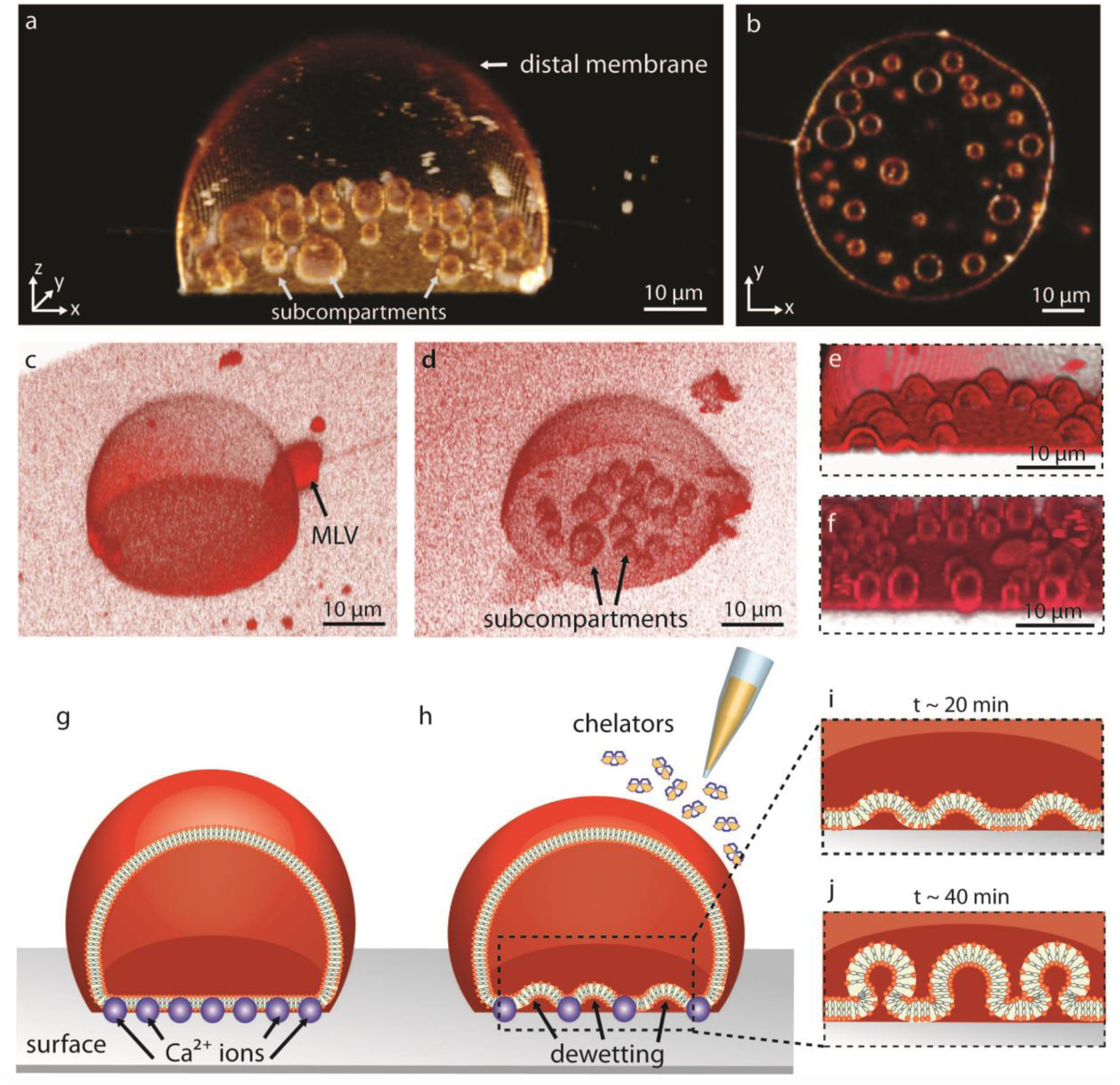
Subcompartmentalization in surface-adhered model protocells. **(a)** Confocal micrograph, reconstructed in 3D, showing a model protocell enveloping several subcompartments. **(b)** model protocell in **(a)** in xy cross sectional view. Confocal micrograph of **(c)** a model protocell adhered on a solid surface in presence of Ca^2+^ **(d)** a model protocell with subcompartments after chelator exposure **(e)** emerging subcompartments ∼20 min. after chelator exposure **(f)** mature subcompartments ∼40 min. after chelator exposure **(g-j)** Schematic representation of the subcompartmentalization process. **(g)** A single model protocell consisting of a single lipid membrane, adheres onto the surface in presence of Ca^2+^. **(h)** Chelators are brought into the vicinity of the compartment, leading to removal of Ca^2+^ followed by subcompartmentalization. The initially surface-adhered bilayer of the model protocell transforms into semi-spherical **(i)**, and later to spherical **(j)** subcompartments. Micrographs in **(c-f)** are from different experiments recorded in identical conditions. Membrane composition of model protocells shown in **(a-f)** is PC-DOPE. The surfaces used in **(a-b)** and **(e-f)** is Al_2_O_3_, and in **(c-d)** is Al. Background fluorescence in **(c-d)** is due to reflection from the Al surface.

### Temperature-enhanced subcompartmentalization

Temperature has a strong influence on membrane viscosity and membrane-surface adhesion^17^. When the temperature was locally increased from 20 °C (room temperature) to 40 °C, the basal membrane started to de-wet the surface, forming subcompartments twice as rapidly. **Fig. 2a-f** show an experiment, during which the subcompartments inside a surface-adhered model protocell form at increased temperature (**Mov. S1**). The subcompartments fuse upon contact and grow in size (**Mov. S1**). For example, the three compartments which are initially distinct (yellow symbols in **panel d**), merge at t=33 min to form a single subcompartment (yellow symbols in **panel f**). A similar experiment is depicted in **Fig. 2g-l** (**Mov. S1**). Both experiments shown in **Fig. 2** have been performed on Al_2_O_3_. The green plot in **Fig. 2m** shows the circumference of the basal membrane depicted in **Fig. 2a-f**, over time. The periphery of the surface-adhered portion of the lipid membrane is not retracting, as the circumference remains constant (**Fig. 2m**, green line). The blue plot in **Fig. 2m** represents the total membrane area utilized by the formation of subcompartments over time. The geometry of the emerging subcompartments has been assumed to be half-spheres in order to estimate the minimally required membrane area. The insets in **Fig. 2m** shows cross sections of model protocells from two different experiments in the xz plane (side view), one at the beginning of an experiment (left) and the other after the subcompartmentalization has occurred (right). The distal membrane of the giant lipid container has decreased in area, indicating that this fraction of the membrane is partially consumed during formation. The plots in **Fig. 2n** show the number (green) and the average diameter (blue points, orange line) of the new compartments in **Fig. 2g-l**. With the emergence and growth of many small subcompartments, the diameter gradually increases over time. Some eventually fuse (inset), intermittently causing sudden drops in the total number (dashed vertical lines in **Fig. 2n**). Subcompartments occasionally collapse (black arrows in **Fig. 2n, Mov. S1**), presumably due to the consumption of their membrane by the competing growth of other containers in their vicinity.

**Figure 2.**
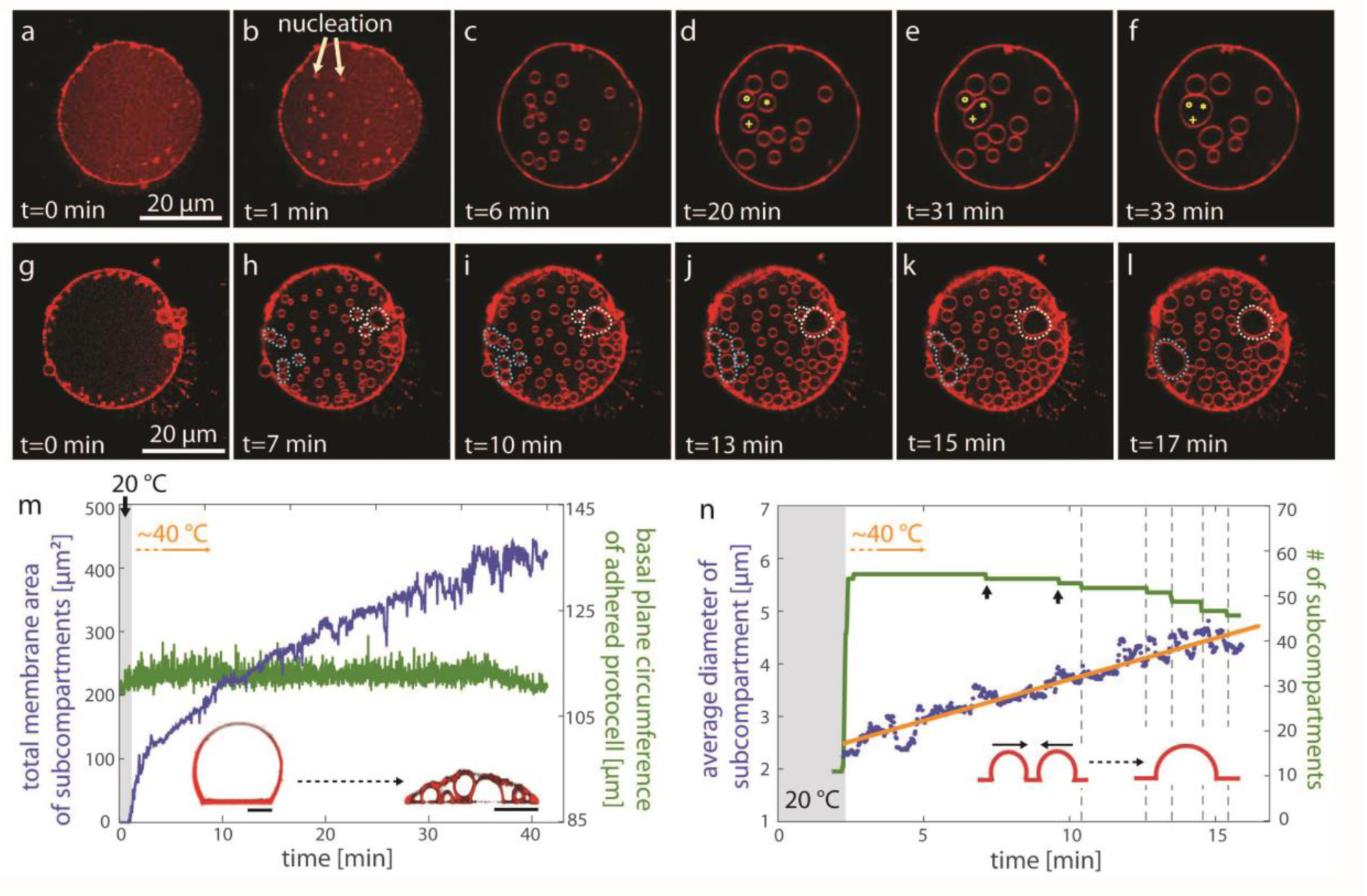
Enhanced subcompartmentalization at increased temperature. **(a-l)** Confocal fluorescence micrographs showing two different experiments **(a-f, g-l)** of temperature-enhanced subcompartmentalization (xy cross sectional view). **(m)** Plots showing the increase in subcompartment membrane area (blue), and the circumference of the protocell (green), *vs*. time, for the experiment shown in **(a-f)**. The inset shows the xz cross sectional view of two protocells from different experiments before and after temperature increase. Scale bars (inset): 10 µm. **(n)** The average diameter and number of subcompartments during the experiment shown in **(g-l)**. The number of subcompartments (green plot) increases instantly with temperature increase, and decreases during fusion or, rarely, due to the sudden collapse of the subcompartments (black arrows). The inset in **(n)** is the schematic drawing of fusion of the subcompartments. The time point of each fusion event is marked with a dashed vertical line. The average diameter (orange line fitted to data points in blue, d=0.0032*t+2.5) gradually increases during subcompartment growth. The membrane of the model protocells shown in this figure is composed of PC-DOPE, and the surface is Al_2_O_3_.

### Encapsulation and isolation of contents

The purpose of subcompartments in a biological cell is the segregation of chemical compounds and processes within individual boundaries inside the main volume of the cell (primary volume). In order to investigate the ability of the subcompartments to encapsulate molecules from the ambient environment and to maintain them, we delivered a water-soluble fluorescent compound to the exterior of each protocell (*cf*. **S2** for the details of the setup) and monitored the transmembrane transfer. **Fig. 3a** shows a 3D micrograph (x-y-z) of a protocell on Al_2_O_3_ with several subcompartments, one of which maintains the encapsulated fluorescein solution. In order to achieve encapsulation in a selected model protocell, the fluorescein buffer (25 µM) was delivered with an open-space microfluidic device^18^(**Fig. 3b**). The presence of the compound can be switched on and off on demand, and the fluorescence signal during the cycles of exposure and removal can be monitored in real time inside the protocell and the subcompartments. We observe that during fluorescein exposure, the subcompartments rapidly encapsulate the compound, and immediately release it upon removal from the surrounding. This indicates that the compartments are not fully closed and have access to the surrounding solution through the space between the surface and the model protocell (**Fig. 3c**). After a certain time point, some subcompartments hold the content and can maintain it during a long period of time, indicating that the size of the opening of the subcompartment is reduced, or completely sealed (**Fig. 3d**). Upon temperature increase, the compartments appear to maintain the content longer.

**Figure 3.**
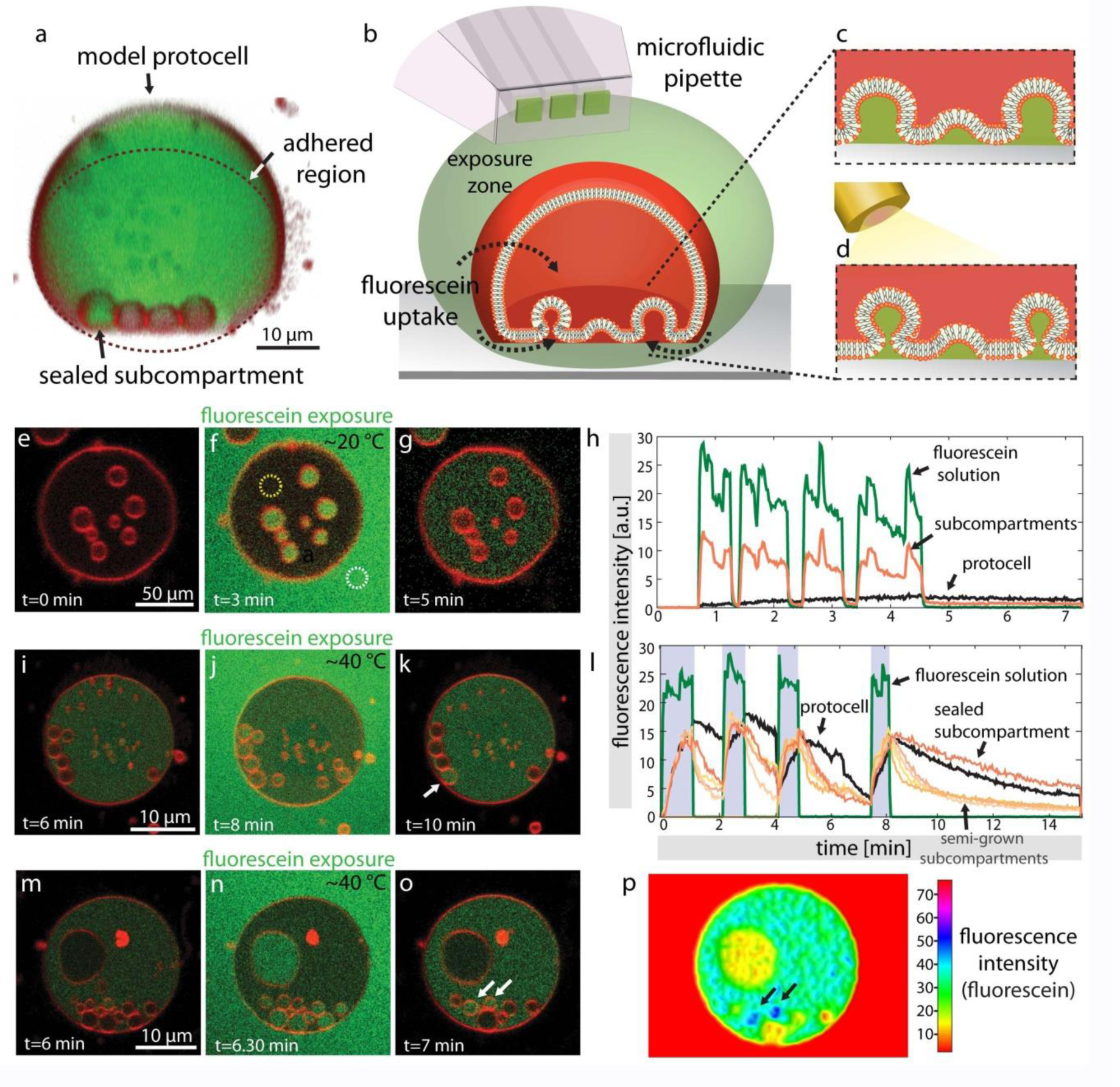
Encapsulation and compartmentalization of fluorescein in model protocells. **(a)** Confocal micrograph of a model protocell, reconstructed in 3D, after pulsing with fluorescein. One of the four subcompartments maintains fluorescein content. **(b-d)** Schematic representation of the fluorescein exposure experiment. **(b)** The microfluidic superfusion pipette creates a fluorescein exposure zone around the model protocell. **(c)** Semi-grown subcompartments encapsulate fluorescein during the exposure through the space at the membrane-surface interface. **(d)** Temperature increase together with fluorescein exposure result in sealed subcompartments encapsulating the fluorescein. **(e-g, i-k, m-o)** Confocal micrographs (xy cross section) showing the fluorescence intensity inside the subcompartments and the primary volume during fluorescein pulsing (***cf*. Mov. S2**). (**h** and **l**) Plots showing the fluorescence intensity of different regions *vs*. time during the experiments shown in **(e-g)** and **(i-k)**, respectively. The temperature increase during **(i-k)** is synchronized with fluorescein pulsing (gray zones in **(l)**). The plots in green color represents the average intensity of the stock fluorescein buffer (circle indicated by white dashed line in **(f)**). The plot in orange color in **(h)** is the arithmetic mean intensity of the subcompartments *vs*. time. The plot in black color in **(h)** is the intensity of the primary volume of the protocell (circle indicated by yellow dashed line in **(f)**). In **(l)** the average intensity of each compartment have been depicted individually (orange-spectrum graphs). The color plot in **(p)** shows the intensity of fluorescein in **(o)**. Black arrows show the position of the two sealed subcompartments corresponding to the entities indicated with white arrows in **(o)**. The sealed compartments display higher fluorescence intensity compared to the other subcompartments and to the primary protocell volume. The membranes in this figure are composed of PE-PG-CA, and the surface is Al_2_O_3_.

**Fig. 3e-g** depicts the time series of an encapsulation experiment at room temperature (*cf*. **Mov. S2**), with the corresponding fluorescein intensity plots of selected regions during multiple exposure cycles (**Fig. 3h**). The fluorescence intensity of three different regions was monitored: the solution outside the protocell (region circled in white dashed lines in **Fig. 3f**), the primary volume of the protocell (**Fig. 3f**, yellow dashed circle) and the intensity of all of the subcompartments, which later was averaged. The plots show an instant encapsulation of fluorescein within the subcompartments with the initiation of the exposure, and immediate release of the contents once the exposure is terminated (**Fig. 3h**). The model protocell, on the other hand, encapsulates the fluorescence molecule gradually and maintains it.

At elevated temperatures (**Fig. 3i-p, Mov. S2**), the leakage of fluorescein from the subcompartments decreases (**Fig. 3l**), compared to the leakage in the experiment depicted in **Fig. 3e-g**, which indicates a reduction of the size of their opening (**Fig. 3k-l and o-p**). The concentration of the fluorescein inside the contracted subcompartment can eventually be slightly higher than in the internal medium of the primary volume (**Fig. 3l** and **3p**).

### Pore-mediated uptake

We then examined the role of transient pores in the uptake of material from the ambient solution. As showed in **Fig. 3**, the fluorescein molecules are encapsulated by the primary volume. To analyze the mechanism of the uptake, we exposed individual surface-adhered protocells without internal compartments to a fluorescein buffer, using an open volume microfluidic device (**Fig. 3, SI section S2**). **Fig. 4a** shows the fluorescence intensity inside 5 different protocells of similar size *vs*. time upon exposure to fluorescein. Uptake rates varying from less than 20% of the stock solution (**Fig. 4a**, plot in pink color) to almost 90% (plot in blue color) were observed. These uptake rates imply that it is not via simple diffusion through the membrane. We attribute this behavior to the presence of transient membrane pores, which has been previously observed and characterized in similar systems^19^. We then compared the experimental values depicted in **Fig. 4a** to the results of finite element model (FEM) simulations shown in **Fig. 4b-d**. The FEM here assumes a single container of size 5 µm with membrane thickness of 1 nm. The model vesicle is exposed to the fluorescein present in the external solution. **Fig. 4c** shows the magnified view of the pore marked with a white frame in **Fig. 4b**.

**Figure 4.**
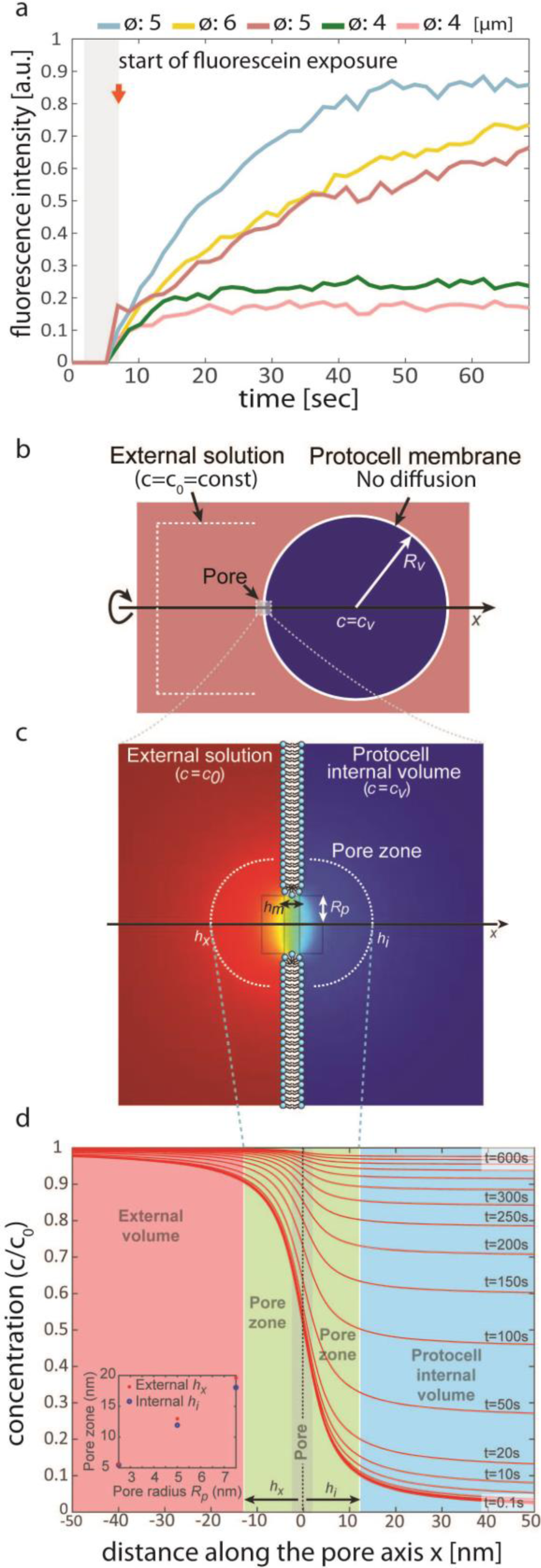
Finite element model (FEM) for uptake through transient nanopores. **(a)** Fluorescein intensity inside the surface-adhered, non-compartmentalized model protocells, over time. **(b-d)** FEM of fluorescein uptake by a model protocell from the external solution through a nano-sized membrane pore. **(b)** A single circular pore (toroidal gap) with a radius of R_p_ = 5 nm is located in the membrane of a giant compartment with radius of R_v_ = 5 µm **(c)** Concentration across the membrane in the vicinity of the pore depicted in **(b)**. The liquid volume near and inside the pore is determined to be the ‘pore zone’ (yellow-green area). **(d)** Concentration profile of fluorescein at the pore zone and inside the protocell at selected time points changing from 100 ms to 600 s. The inset in d shows the relation between the concentration gradient (extent of the pore zone) and the pore radii of 2.5 nm, 3 nm, 5 nm and 7.5 nm. The membrane of the vesicles used in (a) are composed of PC-DOPE; the surface is Al_2_O_3_.

In the model, the passage of molecules is assumed to occur through a single membrane pore (**Fig. 4c**), Ø=5 nm, while the diffusion through the membrane otherwise is prevented. The uptake rates of fluorescein are calculated as the fluorescein diffuses from the external solution into the model protocell through the membrane pore. The diffusion coefficient of fluorescein in water at room temperature, *D* = 4.25*10^−10^ m/s^20^, is taken into consideration. Each plot in **Fig. 4d** represents a different time point upon exposure changing from 0.1 s. to 600 s. The concentration of the fluorescein in the external medium is considered to be 100%. The plots in **Fig. 4d** show final concentrations inside the cell at different time-points, with a steep concentration gradient inside the pore zone (green region). The pore zone is almost symmetrical in both external and internal side of the membrane interface, and reaches a distance of approximately twice the pore radius. The size of the pore zone varies with the changing pore radius (inset to **Fig. 4d**). Further away from the pore, the concentration of the solutes can be considered homogeneous on either side (**Fig. 4d**, red and blue area). The model protocell can have multiple pores further apart with no overlapping pore zones where the transport from each pore contributes linearly. The calculations for different container and pore sizes can be found in **S4**.

### Pseudo-division

In some occasions we observe extensive contraction and rupturing of the distal membrane (**Mov. S3**) which results in disintegration of the original protocell, leaving behind only the intact subcompartments, *i.e*., daughter cells (**Fig. 5**). **Fig. 5a** shows an xz cross section of two subcompartments with the distal membrane in direct contact. **Fig. 5b-c** shows the xz cross section and the perspective view of daughter cells remaining after the original enveloping membrane is disintegrated. The light circular region in **Fig. 5c** consists of a single bilayer, which is also visible in **Fig. 5b**. A double lipid membrane surrounds this region (dark red region in **Fig. 5c**), indicating lipid spreading around an originally isolated intact cell. **Fig. 5d-f** shows the schematic representation of the process in **Fig. 5a-c**. Note that the earlier observations of exchange of material between the primary volume and the ambient buffer on one hand, and the possibility of exchange between the primary volume and the subcompartments on the other hand (**Fig. 5d**), will be discussed in the context of primitive protocell division in the following section.

**Figure 5.**
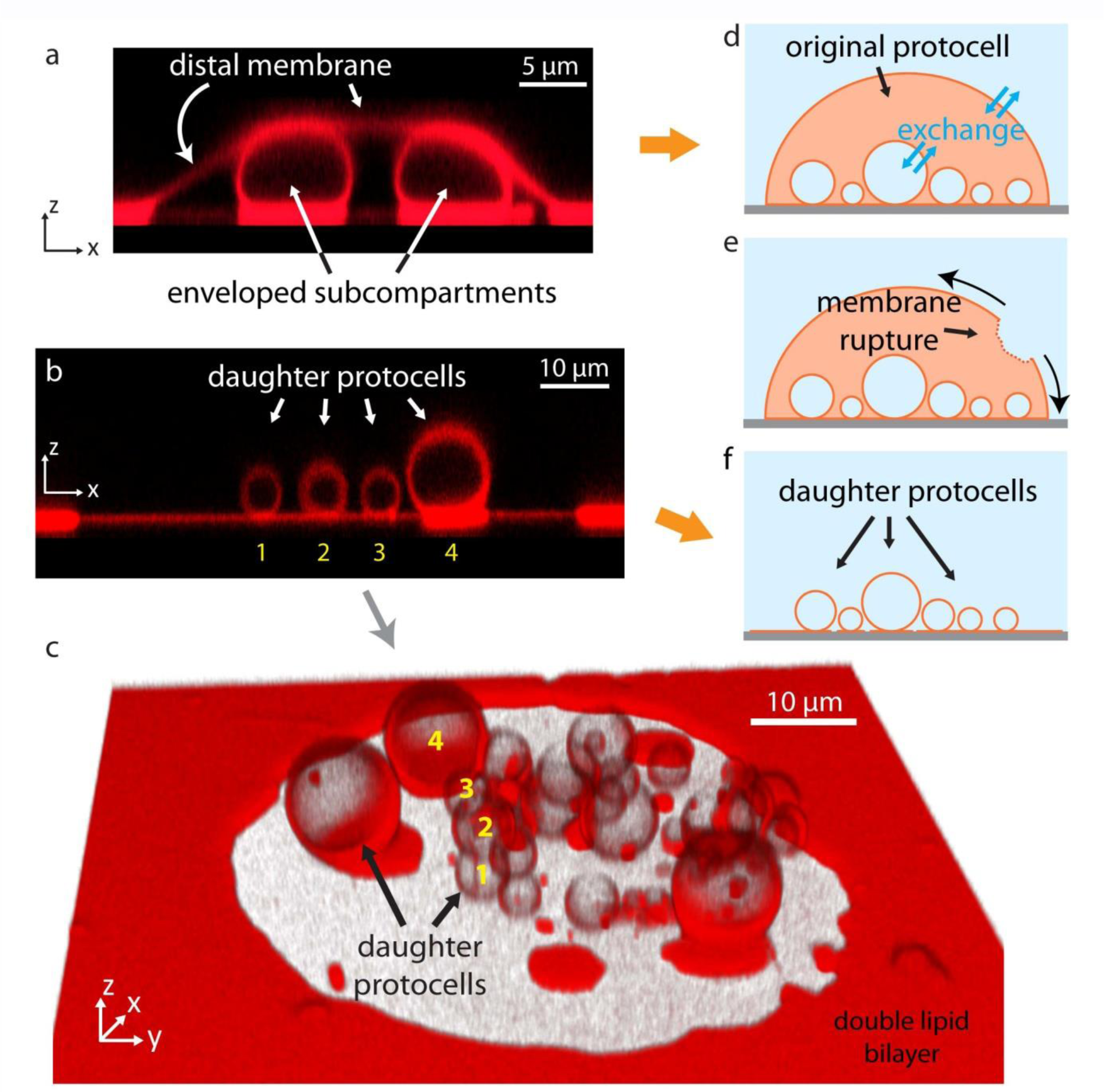
Pseudo-division. **(a)** Confocal micrograph of a subcompartment (xz cross section) with the distal membrane in direct contact. (**b-c**) Confocal micrographs of daughter cells remaining after the original enveloping membrane is disintegrated. (b) xz cross section (c) the perspective view (xyz) **(d)** Light circular region consists of a single bilayer, which is also visible in **(c). (d-f)** Schematic representation of the process in **Fig. 5a-c. (d)** Material between the primary volume and the ambient buffer, and between the primary volume and the subcompartments can be exchanged. The membranes of the vesicles shown in this figure are composed of PC-DOPE; the surface is Al (with a native oxide layer).

## Discussion

### Compartmentalization mechanism

**Fig. 1** shows that the spontaneous compartmentalization occurs when the strength of the adhesion of the protocell to the solid substrate decreases. Adhesion is mediated by the Ca^2+^ ions at the interface of the protocell membrane with the solid surface (depicted by purple spheres in **Fig. 1g-h**). Ca^2+^ ions are known to have a fusogenic effect by binding to the lipid head groups^21^, mediating membrane-surface^15,16,22^, or membrane-membrane pinning^23^. The adhesion of membranes to a solid substrate does not necessarily require Ca^2+^ in the solution^24-26^, although its presence on the surface promotes adhesion. There are several Ca^2+^-rich early Earth minerals, which might have acted as a source^27-29^. The vesicles adhere to the substrate in a dome-like shape^30^. Addition of chelators leads to gradual depletion of Ca^2+^ from the aqueous medium including the space between the surface and the membrane, leading to the removal of pinning points. This causes partial de-wetting at the membrane-surface interface^31^. BAPTA which is one of the two chelators we utilized, has been shown to act as a shuttle buffer, driving Ca^2+^ out of the membrane and decreasing the Ca^2+^ concentration specifically at the membrane interface^32^. Intercalation of EDTA, another chelator we use at mM concentrations, into the lipid head groups has been proposed to cause membrane bending and formation of invaginations^33^. We do not think that direct interactions of BAPTA or EDTA with the membrane is the main cause of the compartmentalization, as in that case the compartments would not only form at the surface but also along the liquid interface to the protocell membrane. Formation of the compartments specifically on the solid substrate is a strong indication that the mechanism is surface adhesion dependent. EDTA and BAPTA are strong metal chelators which were certainly not present on the early Earth. However, clay minerals were abundant in the prebiotic environment which could have played the role of the chelators in our experiments. Clay minerals are known to efficiently adsorb mono- or divalent ions including Ca^2+34^. These phyllosilicates, resemble in surface quality (flatness, homogeneity, surface potential) the synthetic surfaces used in our study^26^. J. Szostak and co-workers showed enhanced vesicle formation upon adsorption of sheets of amphiphiles on various types of solid particle surfaces, among them clays^35,36^.

The creases can also form spontaneously in membranes with low tension^22,37,38^. Once the invaginations form, the transformation of these highly curved membrane regions into larger spherical compartments occurs through a self-driven process, during which the membrane curvature is reduced^39^. A similar study, where the pinning/de-pinning occurs by means of streptavidin-biotin coupling, reports on this phenomenon ^9^. A high membrane curvature in the area of nucleation causes an increase in the membrane tension, which leads to Marangoni flow of lipids from the region of relatively low membrane tension, *i.e*. the distal membrane of the protocell, towards the region of higher tension. The distal membrane of the protocell is consumed as depicted in **Fig. 6** (also in **Fig.1**, inset in **Fig. 2m**). Additionally, an attached MLV (**Fig. 1c**) can serve as a lipid source.

**Figure 6.**
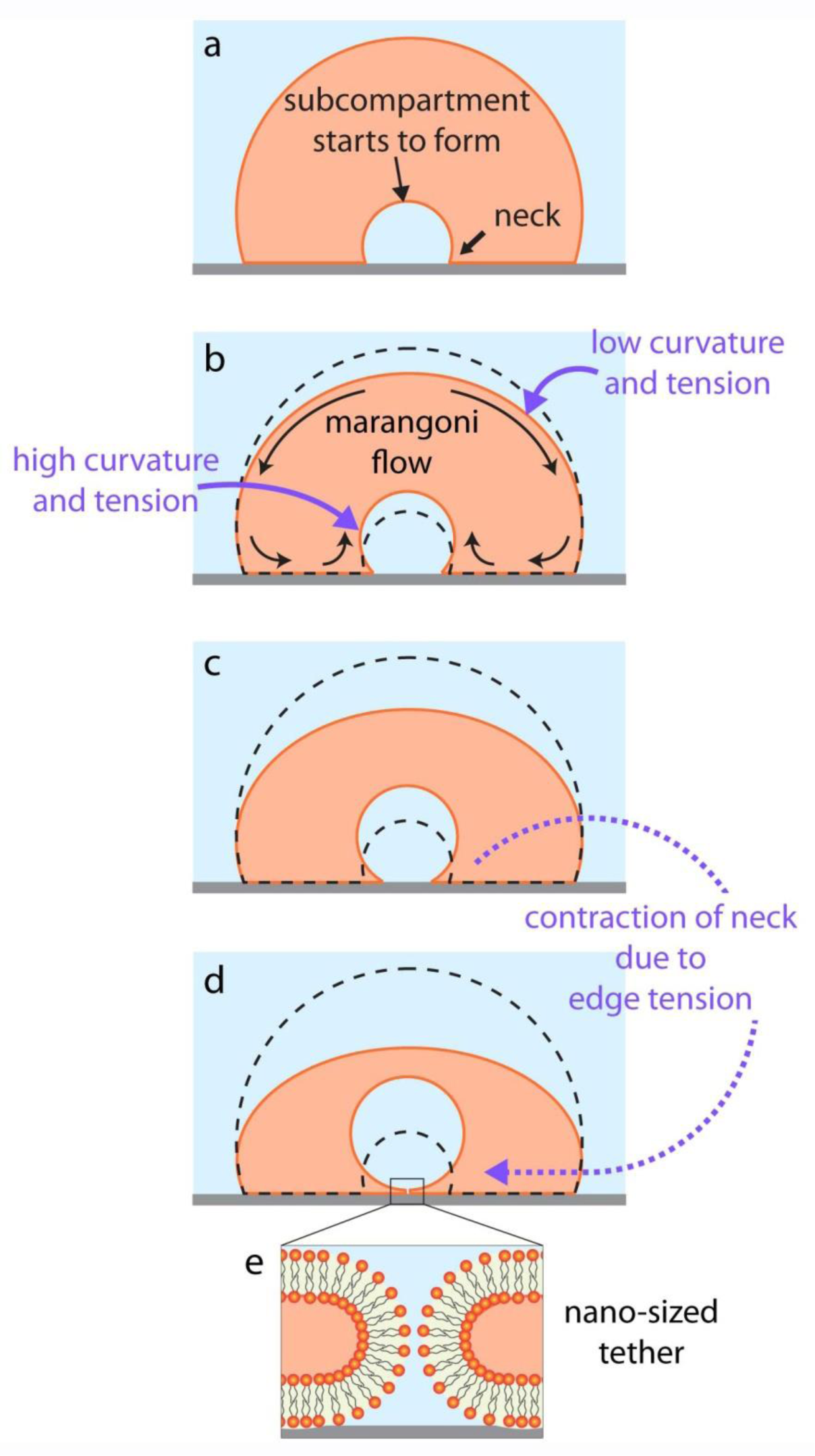
Subcompartmentalization mechanism. **(a-d)** Growth of a subcompartment enveloped in a surface-adhered protocell. During growth the distal membrane is consumed. **(e)** As pinning is removed from the contact line between the newly formed subcompartment and the surface, the contact line will contract due to edge tension transforming to a nano-sized pore or tether. The subcompartment stays connected to the basal membrane through a nano-sized pore or nanotube.

After a subcompartment has matured (**Fig. 6b-d**), it stays connected to the basal membrane through a nano-sized pore-like connection (**Fig. 6e**) as the force requirement for disconnecting the vesicle is very large: in the order of ∼mN^16,40^ *vs*. ∼pN^14^ for pulling a membrane tether that can remain connected through the neck. As pinning is removed from the contact line between the newly formed subcompartment and the surface, the contact line will contract due to edge tension transforming to a nano-sized pore or tether (**Fig. 6e**). The area under the newly formed compartment re-wets even without the adhesion promoter Ca^2+^ lowering the surface free energy of the substrate.

### Temperature-enhanced formation

The role of temperature has been intensely discussed in the origins of life context, in particular whether high temperatures could have been beneficial or detrimental to protocell formation^41,42^. The hydrothermal vent hypothesis which has been criticized for allowing excessively high temperatures, salinity and pH gradients is, after the discovery of “Lost City”-type hydrothermal fields (LCHFs), being reconsidered^42^. LCHFs accommodate temperatures between 40 and 90 °C, which could have promoted the formation of prebiotic membrane compartments^43^.

In our study, temperature increase weakens membrane-surface adhesion and causes immediate surface de-wetting^17^, facilitating the subcompartment formation. Furthermore, it causes the membrane fluidity to increase, and enhances fusion^44,45^. This results in accelerated growth of the subcompartments (**Mov. S1**). While the time scale in the case of subcompartmentalization induced by Ca^2+^ depletion ranges from several tens of minutes to an hour (**Fig. 1**), it only takes a few minutes to form the compartments when the temperature is increased to 40 °C (**Fig. 2a-l**). A nucleated compartment can rapidly mature and be internalized by the protocell (**Fig. 2m, inset**) where some of them coalesce upon physical contact. A recent study has reported on the merging of lipid vesicles if they are in close proximity in a locally heated zone^46^. Note that in order to elevate the temperature locally under avoidance of strong convection, we used an optical fiber and IR-B radiation (1470 nm, ***cf*. S2** for the details of the setup).

### Encapsulation and compartmentalization of external compounds

Confining the reactants inside a membrane enclosed space and thereby bringing them into the close proximity can enhance reaction rates^47^ which is a benefit of compartmentalization in modern cells. We used anionic fluorescein as a model molecule to identify the mechanism of compartmentalization and encapsulation events. We have observed the internalization of fluorescein molecules inside the semi-grown subcompartments, delivered locally by superfusion with a microfluidic pipette, by laser scanning confocal microscopy (**Fig. 3**). A negatively charged dye was used to avoid electrostatic charge-based adsorption of the dye to the negatively-charged membrane^48^. When the protocell is repeatedly pulsed with fluorescein, the fluorescence intensity inside the compartments instantly follows the external aqueous environment (**Fig. 3e-h, Mov. S2**). The compartments take up fluorescein which immediately leaks out when the superfusion is terminated. This indicates that the subcompartments encapsulate the compound through the open interface to the surface, which exposes the compartments’ interior to the external aqueous space. Encapsulation of the liquid at a membrane-surface interface by the invaginations has been previously reported. In that system, a protein-ligand coupling has been employed to promote the formation of the invaginations^9^. If the subcompartment encapsulating the compound is suddenly sealed or its opening significantly reduced in size, the instant leakage of fluorescein is prevented and the fluorescence intensity decreases slowly (**Fig. 3l**). While the leakage from the subcompartments is rapid in the initial superfusion cycles throughout the experiment, after t=∼8min the leakage from one particular subcompartment slows down (dark orange plot). Most importantly, the leakage from the subcompartments induced by heating is considerably slower, compared to the Ca^2+^ depletion-induced case (**Fig. 3l** *vs*. **Fig. 3h**). Higher temperatures increase membrane fluidity, causing faster compartment growth. The compartments mature faster, likely resulting in contraction of the opening at the surface interface (**Fig. 3c-d, Fig. 6**) to a nanosize pore or a nanotube (**Fig. 6e**) and, accordingly, prolonged containment of internalized content **(Fig 3i-l)**. This is in agreement with the finite element model. A single nanopore allows the equilibration of material in 600 s. (**Fig. 4**, *cf*. FEM discussion below). This is confirmed by earlier studies on fluorescein diffusion through nanosized conduits^49^.

### Evidence for pore-mediated encapsulation in the primary volume

The primary volume of the model protocell encapsulates fluorescein during multiple cycles of fluorescein exposure (**Fig. 3**). We performed identical superfusion experiments on model protocells without subcompartments, *i.e*. without the involvement of chelators. For different model protocells of comparable size under identical exposure conditions, the amount of encapsulated fluorescein at a given time varies from less than 20% to almost 90% of the concentration in the external medium (**Fig. 4a**). This indicates that the means of transport through the membrane is not diffusion, which would result in each experiment in similar concentrations of the encapsulated compound at a given time. Our results are consistent with the presence of transient nano-sized pores in the membrane through which the fluorescein is diffusively internalized. It is well established that osmotic swelling of lipid vesicles causes an increase in membrane tension which leads to the formation of transient pores in the membrane ^19,22^. The adhesion of protocells to the substrate can cause a similar tension increase^16,22^. The finite element method (FEM) simulations in **Fig. 4b-d** shows that transport through a single or a few nano-sized pores in the lipid membrane of a container of 5 µm size can lead to similar fluorescein concentrations inside the protocells shown in **Fig. 4a**. According to the FEM simulations, during 100 s of uptake through a single pore, the concentration of the encapsulated fluorescein inside the protocell reaches 50% of the concentration of fluorescein in the external medium (**Fig. 4d**). In one of the experiments (red plot in **Fig. 4a**), the internal volume reaches 50% of the concentration of the stock fluorescein solution in 20-30 s. This corresponds to 3-5 membrane pores. It is reasonable that since 3-5 pores of nm size belong to a protocell of micron size, the pores would not overlap, and the pores would not jeopardize the integrity of the vesicle.

### Implications for the origins of life

Origin of life studies often focus on a compartment as a whole - the (model) protocell. The possibility of consistent formation of organelle-like structures inside a protocell has not been considered plausible due to the lack of proteins and membrane-shaping machinery, which would transform the original membrane material to modular subcompartments and further stabilize them^10^. Solid surfaces, which were abundant on the early Earth in the form of minerals, could to some extent have mimicked the role of sophisticated machinery. Surfaces intrinsically possess energy, which can promote membrane-surface interactions in several different ways^26^. Previous studies have reported on the enhancing effects of surface associated bio- and organic chemistry relevant to the *RNA world*^13^, including the synthesis of prebiotic peptides^12^, nitrogen reduction, lipid self-organization, condensation-polymerization reactions, selection and concentration of amino acids and sugars as well as chiral selection^50^. The reported surface-dependent phenomenon might pose an advantage over multi-vesiculation observed at the compartment membranes in bulk solution^51^. Mineral surfaces have recently been shown to also induce protocell formation^35,36,39^. Our findings reveal the capabilities of high energy surfaces, which are able to induce subcompartmentalization of adherent model protocells. We selected components which have direct representations in the prebiotic environment on the early Earth.

### Flat solid oxide substrates as mineral models

The solid model substrates Al_2_O_3_ and Al (with a native oxide layer) as well as SiO_2_, represent constituents of rock forming minerals on the early Earth^52^. Al_2_O_3_ occurs naturally as Corundum, a mineral which has been studied and shown to amplify lipid bilayer formation^52^. Sputtered SiO_2_ corresponds to Quartz, which was associated with amino acid adsorption and peptide synthesis^13,53^.

### Protocell models

We used giant unilamellar vesicles as protocell models. They consist of phospholipid molecules which are part of the origins of life discussion. Several types of phospholipids have been synthesized in the laboratory under prebiotic conditions^54-57^. Their precursor molecules were found in fluid inclusions in minerals relevant to the early Earth^53^. A detailed list of the lipid membrane compositions and the surfaces leading to phenomena we describe in this study is provided in **S1**.

### Pseudo-division

**Fig. 3l** reveals similar intensity levels of encapsulated fluorescence in the primary volume of the model protocell, and in one of its subcompartments. It is conceivable that the encapsulated contents are transported between the subcompartment and the primary volume through transient nano-sized membrane pores. We assume that the pores are present in the protocell membrane, which we have experimentally verified. Since the membrane surrounding the primary volume and the subcompartments are connected, and pores/defects are mobile in a fluid membrane, it is reasonable to assume that the pores can also be present in the subcompartment membrane. This would allow the transport of material across the membrane from the primary volume to the sub compartment. In some occasions the distal membrane of the model protocell ruptures, leaving multiple subcompartments, *i.e*. the daughter cells, exposed directly to the bulk solution (**Fig. 5**). This can be perceived as pseudo-division as this is not a direct splitting of the initial compartment, but a two step process starting with the transfer of internalized material to the daughter cells, followed by the disappearance of the mother protocell. We suggest that the surface mediated subcompartmentalization can therefore be viewed as starting point (pre-division) for protocell replication and division.

## Conclusion

The question of how early in evolution cells could have had membranous subcompartments came recently into focus after their existence in archaea and bacteria has been shown. In this study we report that membrane-enveloped subcompartments consistently form in model protocells adhered on mineral-like solid substrates. The formation requires a minimum set of essential components: an amphiphile compartment, a solid surface and an aqueous environment. External compounds can be encapsulated and transported between the primary volume of the protocell and the subcompartments. We show that when a protocell disintegrates, the subcompartments remain intact, and adhered to the substrate, suggesting the possibility of a primitive form of division. Following the earlier findings on enhancing effects of surfaces in prebiotic chemistry, this advocates that protocells with membraneous subunits simultaneously running multiple reactions could have existed at the origin of life. If the process can be repeated in cycles over several protocell-subcompartment generations remains an interesting question to be elucidated.

## Materials and Methods

### Preparation of lipid vesicles

To prepare the lipid suspensions, the dehydration and rehydration method^14,58^ was used. Briefly, lipids (99 wt %) and lipid-conjugated fluorophores (1 wt %) (for a detailed list of lipid compositions *cf*. **S1 Table**) were dissolved in chloroform leading to a final concentration of 10 mg/ml. 300 µl of this mixture was then transferred to a 10 ml round bottom flask and the solvent was removed in a rotary evaporator at reduced pressure (20 kPa) for 6 hours to form a dry lipid film. The film was rehydrated with 3 ml of PBS buffer (5 mM Trizma Base, 30 mM K_3_PO_4_, 3 mM MgSO_4_.7H_2_O, 0.5 mM Na_2_EDTA, pH 7.4 adjusted with 1 M H_3_PO_4_) and stored at +4 °C overnight to allow the lipid cake to swell. The sample was then sonicated for 25 s at room temperature, leading to the formation of giant compartments with varying lamellarity. For sample preparation, 4 µl of the resulting lipid suspension was desiccated for 20 min and the dry residue rehydrated with 0.5 ml of 10 mM HEPES buffer containing 100 mM NaCl (pH 7.8, adjusted with 5 M NaOH) for 5 min. The lipid suspension was subsequently transferred onto a solid surface submerged in 10 mM HEPES buffer with the formulation mentioned above, but with the addition of 4 mM CaCl_2_.

### Surface preparation

SiO_2,_ Al and Al_2_O_3_ surfaces were fabricated in the Norwegian Micro- and Nano-Fabrication Facility at the University of Oslo (MiNaLab). All thin films were deposited on glass cover slips (Menzel Gläss #1, 100-150 µm thickness; WillCo Wells B.V., Amsterdam, NL). No pre-cleaning was performed on glass substrates before deposition. SiO_2_ and Al were deposited onto the glass substrates by E-beam and thermal PVD evaporation using an EvoVac instrument (Ångstrom Engineering, Canada), to a final thicknesses of 10 nm of Al, and 84 nm of SiO_2_. Al_2_O_3_ was deposited onto glass substrates by atomic layer deposition (Beneq, Finland), to a final thickness of 10 nm. Surfaces were used immediately after their fabrication.

### Addition of chelators and fluorescein molecules

For initiation of the subcompartmentalization, ambient buffer in the sample was gently exchanged with 10 mM HEPES buffer containing 100 mM NaCl, 10 mM EDTA and 7 mM BAPTA (pH 7.8, adjusted with 5 mM NaOH), using an automatic pipette 20 min after the initial deposition of the vesicles onto the substrates.

The experiments involving fluorescein exposure were performed by using a microfluidic pipette (Fluicell AB, Sweden). The surface-adhered vesicles were superfused with HEPES buffer with 100 mM NaCl, containing 25 µM of Fluorescein Sodium Salt (pH 7.8, adjusted with 5 mM NaOH). The HEPES buffer used in the experiments shown in **Fig. 4a** contains Ca^2+^, but is chelator-free. The vesicles used for **Fig. 4a** are composed of identical lipid species, and are of similar size (4-6 µm in diameter).

### Local heating

An optical fiber coupled to an IR-B laser (*cf*. **S3**) was assembled to locally increase the temperature in the sample^46,59^. A semiconductor diode laser (HHF-1470-6-95, λ = 1470 nm, Seminex), driven with an 8 A power source (4308 Laser Source, Arroyo Instruments) was used in combination with an 0.22 NA multimode optical fiber with 50 µm core diameter (Ocean Optics). The fiber was located around approx. 30-50 µm from the vesicle. The laser current utilized for experiments were in the range 0.7 A, resulting in a local temperature increase to 40 °C^46^.

### Microscopy imaging

All microscopy images were acquired with a laser scanning confocal microscopy system (Leica SP8, Germany) using a HCX PL APO CS 40x oil, NA 1.3 objective. The excitation/emission wavelengths varied with the employed fluorophores: Rhodamine ex: 560 nm/em: 583 nm (**Fig. S1**); Texas Red DHPE ex: 595 nm/em: 615 nm (**Fig. 1, Fig. 2, Fig. 4, Fig. 5**); ATTO 655-DHPE ex: 655/em: 680 (Fig. 3); Fluorescein ex: 488 nm, em: 515 nm (**Fig. 3, Fig.4**).

### Image analyses

3D fluorescence micrographs were reconstructed in Leica Application Suite X Software (Leica Microsystems, Germany). Image enhancement of fluorescence micrographs for the figures were performed with the Adobe Photoshop CS4 (Adobe Systems, USA). The fluorescence intensity analyses shown in **Fig. 2m-n, Fig. 3h, Fig. 3l, Fig. 3p** and **Fig. 4a**, were performed by using NIH Image-J Software. For the micrograph series represented in **Fig. 2m-n (Mov. S1)**, median filtering was applied with Image-J. All graphs were plotted in Matlab R2018a, which was also used to generate the linear fit in **Fig. 2n**. Schematic drawings were created with Adobe Illustrator CS4 (Adobe Systems, USA).

### Finite-element simulations

Finite element modeling (FEM) was performed with the COMSOL Multiphysics software, using transport of dilute species physics (chds). Our model assumes a membrane thickness of *h*_m_ = 1 nm, and fluorescein diffusion coefficient of *D* = 4.25*10^−10^ m/s^20^. The geometry was built using cylindrical symmetry around axis *x*. The vesicle has a spherical shape with radius *R*_v_, which was set to 2.5 μm, 5 μm or 7.5 μm. We assumed no material transport through the membrane (no flux boundary) except through one cylindrical pore (length *h*_m_ and radius *R*_p_) positioned on the *x*-axis and connecting the vesicle with the external volume. The vesicle had initial internal concentration *c* = 0, while external volume had *c* = *c*_0_ = 1. The outer boundary of the external volume was set to the constant concentration 1, mimicking a very large bath compared to the vesicle. For the simulation we varied the vesicle dimension, while the pore dimension was set to *R*_p_ = 5 nm, or changed to a pore radius between 2.5 nm and 7.5 nm while the vesicle radius was kept *R*_v_ = 5 μm (*cf*. **S4**). The simulation was transient from 0 to 600 s in 100 ms steps.

## Supporting information

Spustova_etal_SI

## Acknowledgements

This work was made possible through financial support obtained from the Research Council of Norway (Forskningsrådet), Project Grant 274433, UiO: Life Sciences Convergence Environment, the Swedish Research Council (Vetenskapsrådet), Project Grant 2015-04561, as well as the startup funding provided by the Centre for Molecular Medicine Norway (RCN 187615), and the Faculty of Mathematics and Natural Sciences at the University of Oslo. We thank Dr. A. Jesorka from the Biophysical Technology Laboratory at Chalmers University of Technology, Sweden, for support with the microheating setup, and for stimulating discussions.

## Author Contributions

I.G. and A.A. designed the study, E.K. made the initial setup. K.S. carried out the microscopy experiments and analyzed the experimental data. A.A. developed the FEM model. I.G. suggested the investigation of the surface-induced subcompartmentalization phenomena, contributed to data evaluation, and supervised the project. All authors contributed to the writing of the manuscript.

## Conflict of Interest Statement

AA is a co-inventor of the multifunctional pipette, and minority share holder of Fluicell AB, the company that markets the multifunctional pipette. No payments or financial gain were a reason for, or a direct consequence of, the research contained within the manuscript.

