## Supplementary material for "Subcompartmentalization and pseudo-division of model protocells": Spustova_etal_SI

**Table of contents**

S1 Lipid compositions and surfaces

S2 Experimental setup

S3 Heating system

S4 Finite element simulations

### S1. Lipid compositions and surfaces

#### Table S1. Lipids, lipid-conjugated fluorophores and solid surface combinations

| Surface | Lipid species | *wt%* percentage | Fluorophore (1%) | Associated figure |
|---|---|---|---|---|
| $Al_2O_3$ | PC-DOPE | 69:30 | Texas Red-DHPE<br>ATTO 655-DHPE | Fig. 1a-b, e-f<br>Fig. 2a-l<br>Fig. 4a |
| | PE-PG-CA | 67:23:9 | 16:0 Liss Rhod PE | Fig. 3a, e-p |
| Al | PC-DOPE | 69:30 | Texas Red-DHPE | Fig. 1c-d<br>Fig. 5a-c |
| | PE-PG-CA | 67:23:9 | 16:0 Liss Rhod PE | Fig. S1a |
| $SiO_2$ | PE-PG-CA | 67:23:9 | 16:0 Liss Rhod PE<br>ATTO 655-DHPE | Fig. S1b |

Abbreviations used in the table are as following:

PC - Phosphatidylcholine
DOPE – 18:1 Dioleoylphosphoethanolamine
PE - Phosphoethanolamine
PG - Phosphatidylglycerol
CA – Cardiolipin

All lipid products and 16:0 Liss Rhodamine PE were purchased from Avanti Polar Lipids, USA. Texas Red-DHPE was purchased from Sigma-Aldrich, USA. ATTO 655-DHPE was purchased from ATTO-TEC, Gmbh Germany.

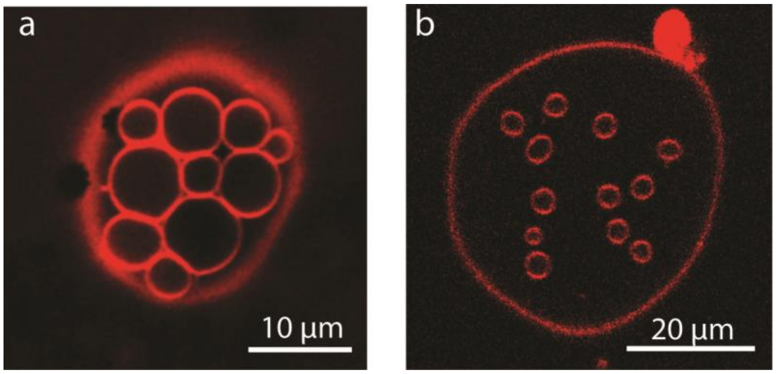


**Figure S1**. Subcompartmentalized model protocells made from PE-PG-CA lipids on (a) an Al surface with a native oxide layer, (b) $SiO_2$ surface.

### S2. Experimental setup

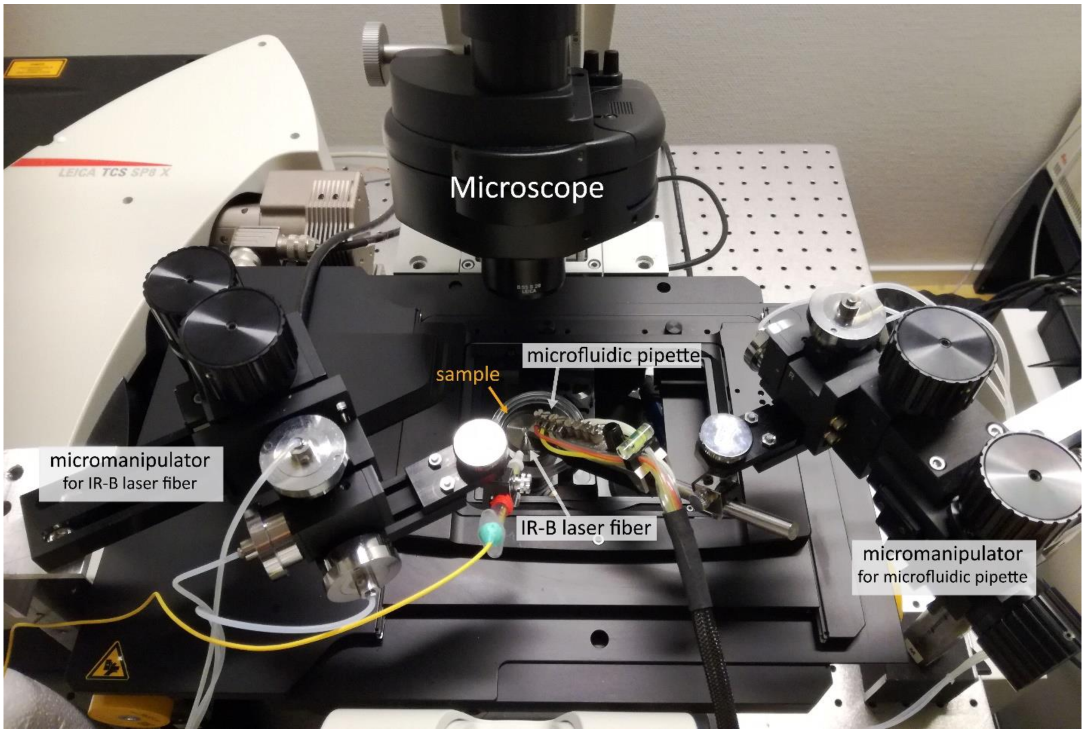


**Figure S2**. For the simultaneous heating and pulsing of fluorescein, an optical fiber coupled to an IR-B laser and a microfluidic pipette were positioned above the model protocells in the sample chamber, using 3-axis water hydraulic micromanipulators. The tip of the fiber and the tip of the pipette were placed in close proximity to target the same model protocell.

### S3. Localized heating

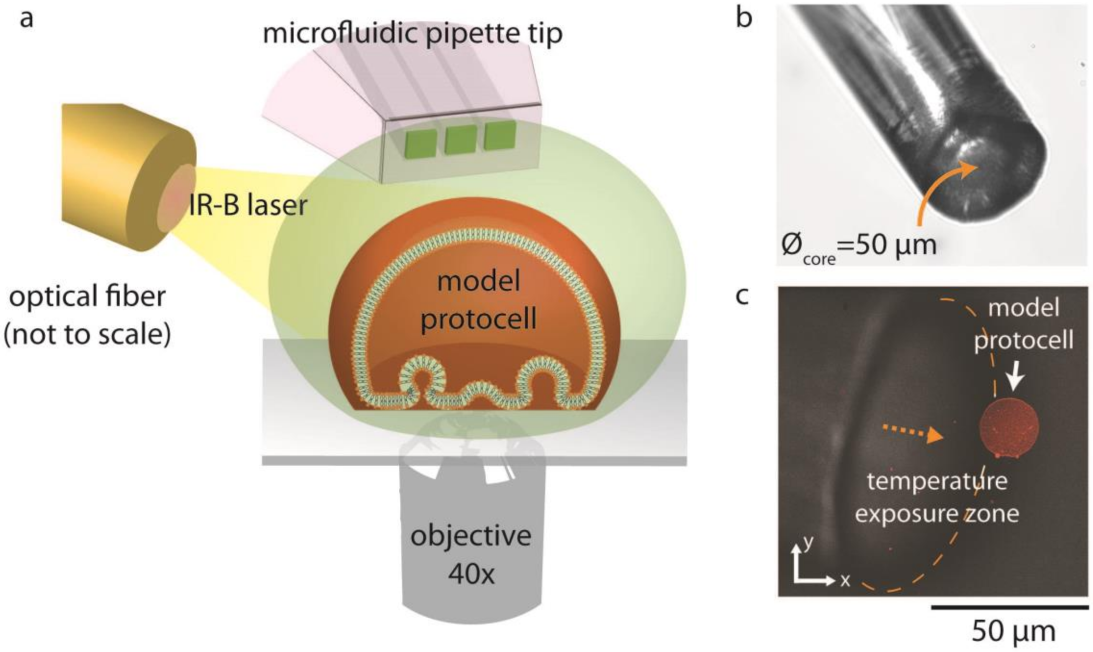

**Figure S3**. Localized heating. **(a)** The optical fiber is positioned above the surface near the model protocell of interest. Upon activation of the laser, the temperature in the vicinity of the protocell is increased to 40 °C. The heating can be applied with (**Fig. 3**) and without (**Fig. 2**) simultaneous fluorescein pulsing. Both the optical fiber tip and the microfluidic pipette tip are submerged into the sample buffer (not shown). **(b)** Micro-photograph of the optical fiber tip employed in the experiments. **(c)** Close up of the optical fiber tip while heating a model protocell.

### S4. FEM simulations

**Fig. S4** shows the concentration change in vesicles of varying size and with varying pore sizes over time, according to the FEM simulations described in the main article (**Fig. S4a**). Data sets are fitted with the function $1\text{-}e^{-kt}$, where the $k$ is a loading rate. This is analogous to charging a capacitor through a resistor (RC circuit), where the vesicle internal volume will take the place of capacitor, and the pore will be the resistance for the loading. As expected, the loading rate increases with pore radius (**Fig. S4b**) and it is inversely proportional to the vesicle volume (**Fig. S4c**).

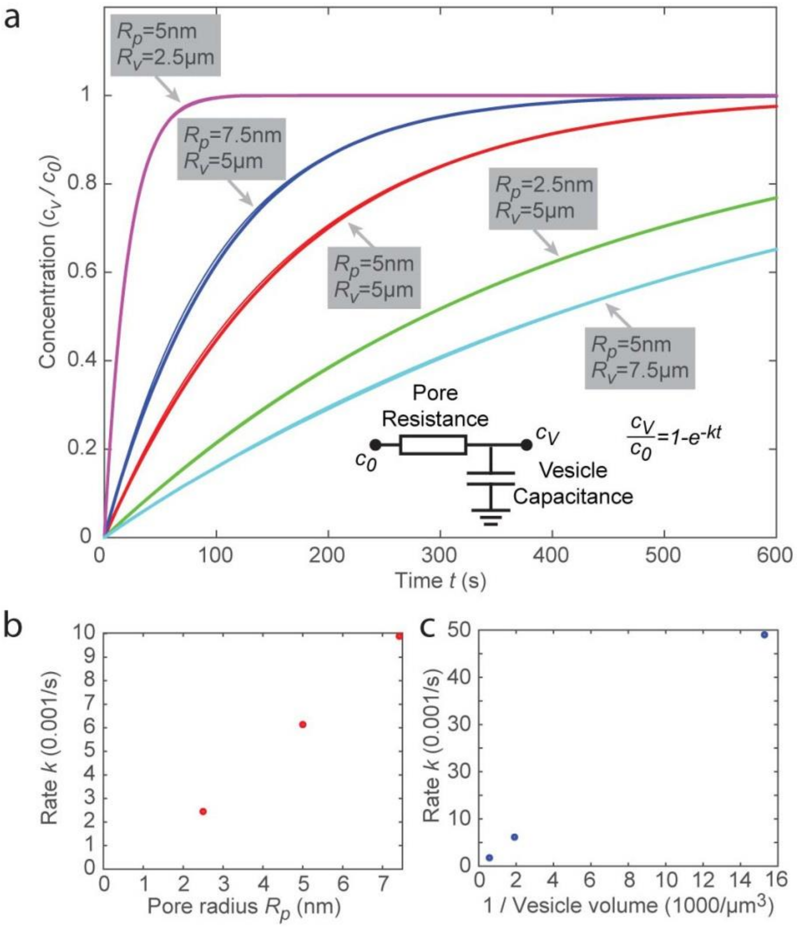


**Figure S4**. **(a)** FEM simulations showing encapsulation of fluorescein in vesicles of varying size and pore size, over time. **(b)** loading rate vs. pore radius, **(c)** loading rate vs. vesicle volume.
